## Supporting Material Figure S1 for "Fast sequence-based microsatellite genotyping development workflow"

Supporting Figure S1 : Effect of sequence coverage (usuing one, two or three Illumina Miseq nano flowcells) on SSRseq data quality for *Alosa* species.

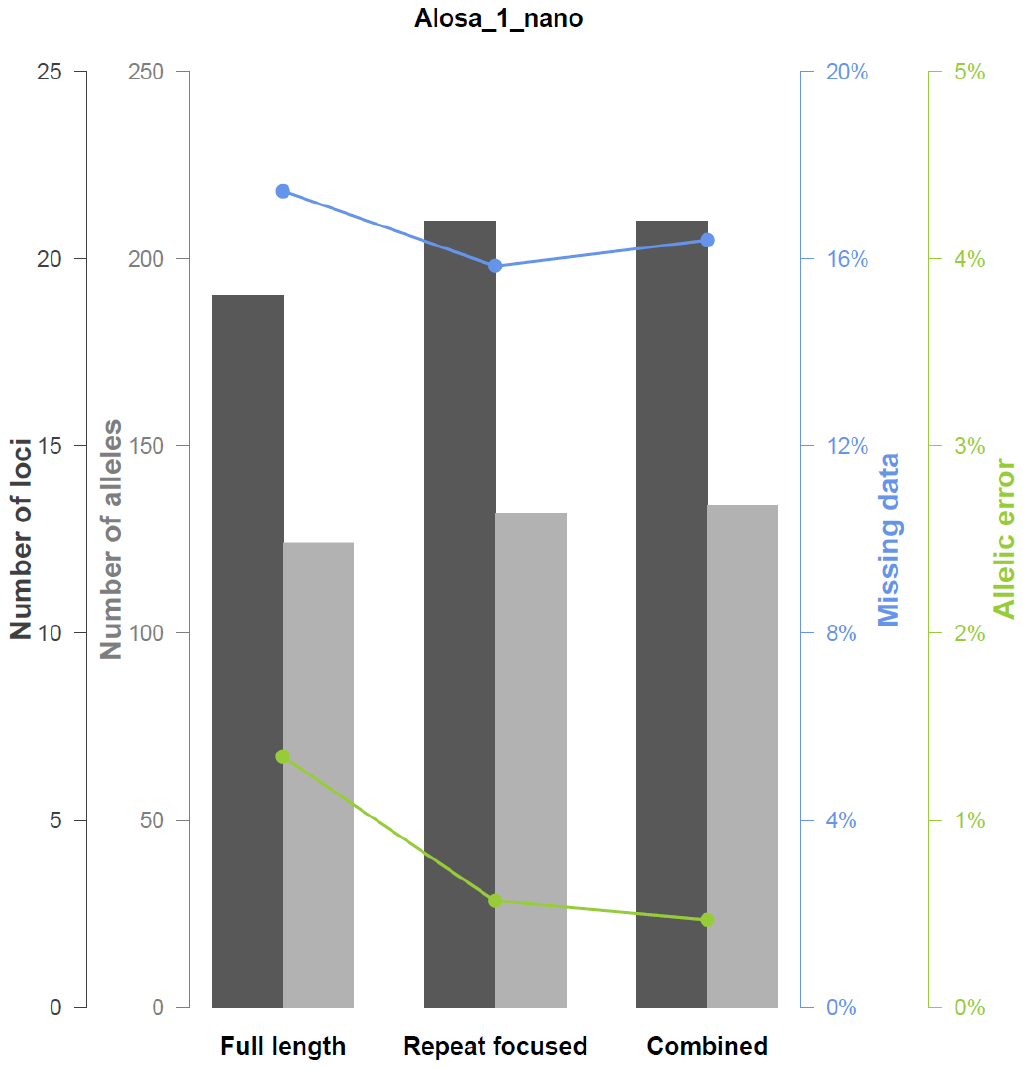

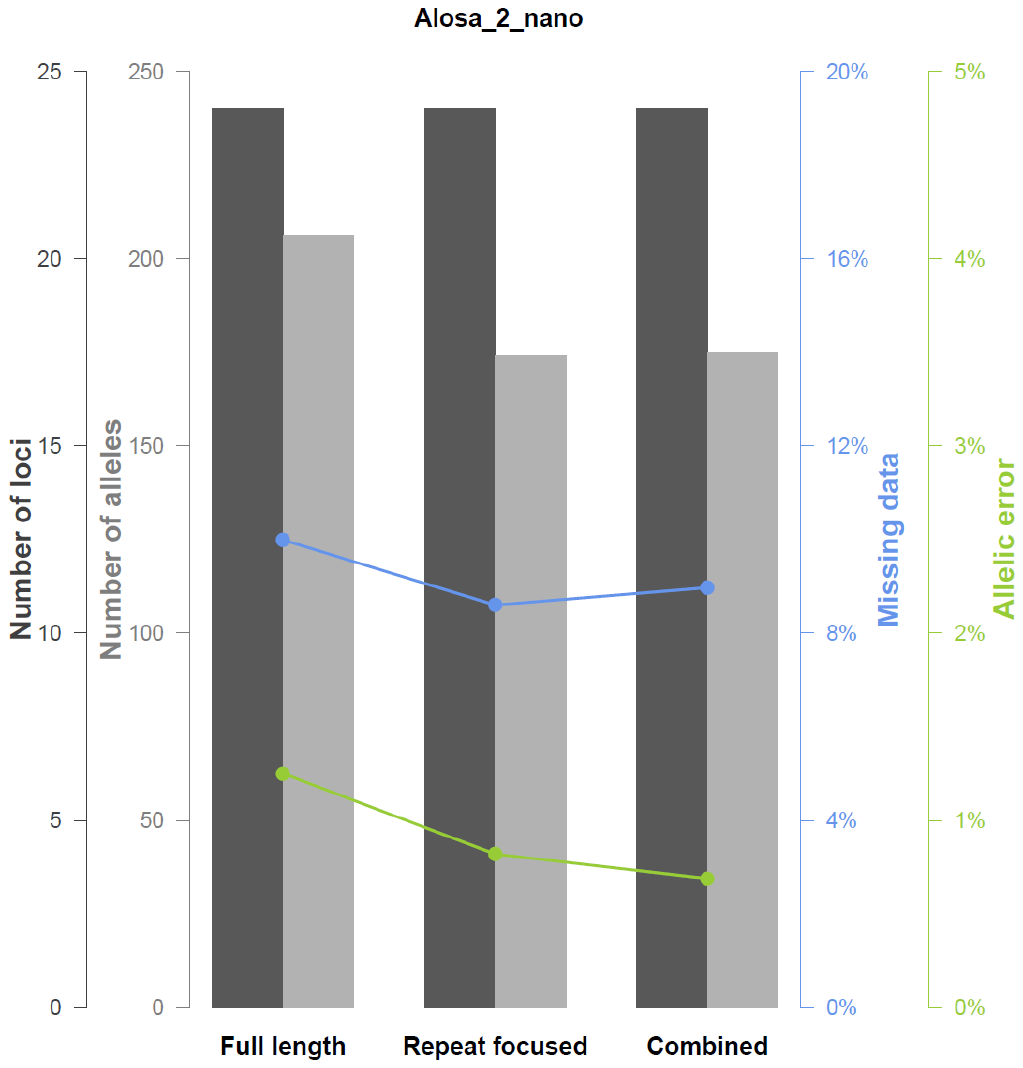

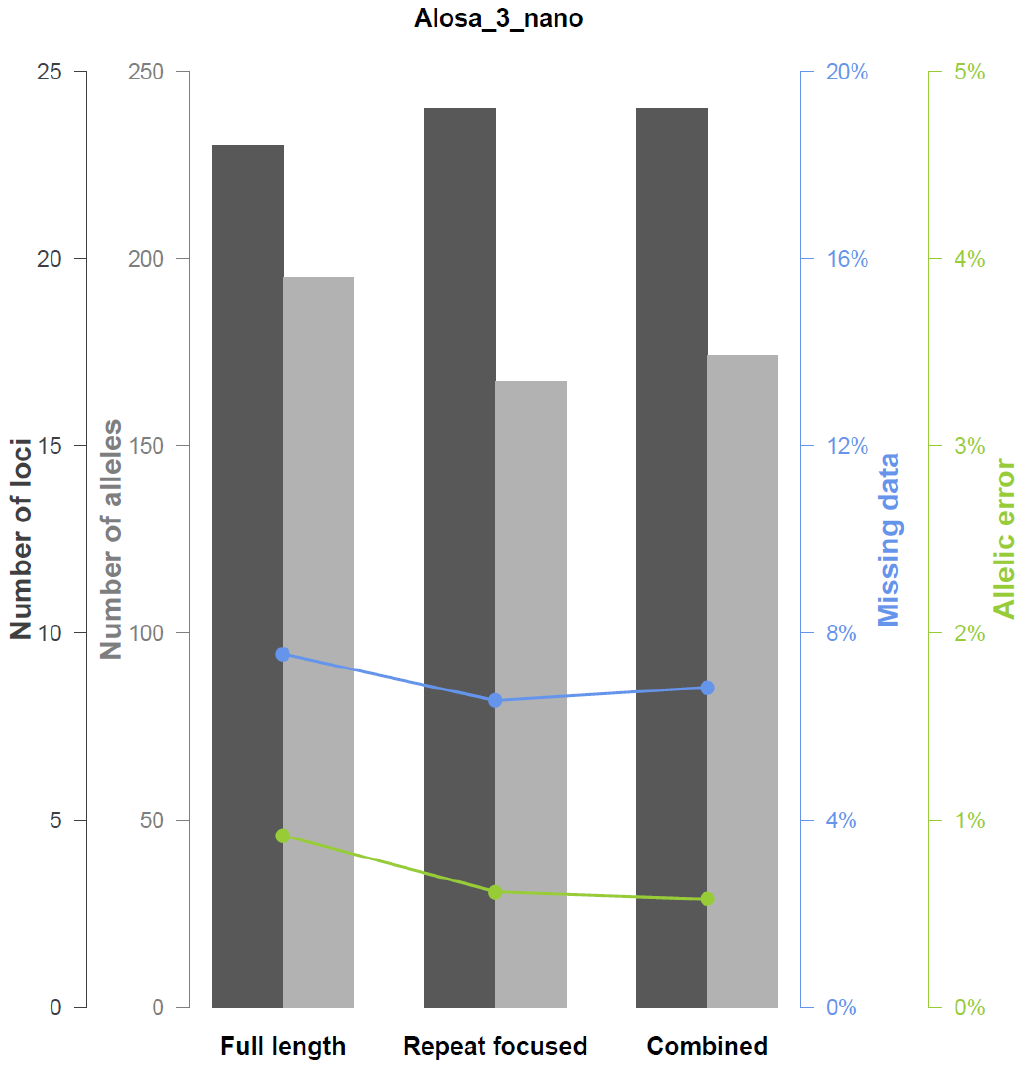
