## Supporting Materials and Methods for "Fast sequence-based microsatellite genotyping development workflow"

### Supporting Materials & Methods

#### Detailed description of FDSTools parameter sets used (PS1 and PS2)

Preliminary analyses showed that no single set of pipeline parameters resulted in good quality genotypic call across all locus. Instead, specific sets of parameters must be used to optimise genotypic call quality for each locus. However, testing numerous parameter combinations would rapidly become cumbersome and impractical. We thus restricted our analysis to two sets of parameters, called PS1 and PS2 thereafter. PS1 and PS2 were selected on the basis of high quality selection for locus showing moderate to high stutters and unbalanced allele coverage for heterozygote genotypes, because such patterns were typical of the loci developed here in non-model species. For Stuttermark, a sequence with a loss or a gain of one repeat compared to another sequence but with high coverage will not be flagged as potential stutter, depending on user defined thresholds controlling for the relative coverage of the two sequences (parameter –s). For PS1, we assumed that a sequence in stutter configuration with -1 repeat is flagged as stutter if its coverage is below 50% of the originating sequence. For PS2, we assumed that a sequence in stutter configuration with -1 repeat is flagged as stutter if its coverage is below 70% of the originating sequence. For -2 stutter sequence, the rule is set to 50% and 70% for PS1 and PS2 respectively when comparing -2 and -1 repeated sequence. In both PS1 and PS2, a sequence showing a gain of one repeat is flagged as stutter if the coverage if lower than 10% of the originating sequence. In all cases, sequence with a coverage of one read for any individuals and locus is not evaluated by Stuttermark and flag as noise (parameter –m 2). Allelefinder ignores sequences flagged as noise or stutter and call one or two alleles from all potential allele based on user defined coverage thresholds. For PS1, a heterozygote genotype is called if the second allele has at least 15% of coverage compared to the highest covered allele for a locus (parameter –m 15). For PS2, this parameter was set to 10% (-m 10). PS1 and PS2 shared some parameter settings: we required a minimum of 20 reads for the most covered allele for a locus in an individual (parameter –n 20), otherwise a missing genotype is called. In addition, specific parameter controlled for potential sample contamination were adjust in accordance to the other parameters: a locus is called missing data if the most covered non-allelic sequence is at least 75% of the most covered allele (parameter –M 75) and a sample is rejected if more than 10 loci have a high number of non-allelic sequences (parameter –x 10). A single execution of the custom-made bash script (SSRseq_DataAnalysis_ParametersComparison.sh available at https://doi.org/10.15454/HBXKVA) run the analysis using the two parameter sets PS1 and PS2 for direct comparison of locus-specific genotypic call quality and selection of the most appropriate settings for each locus.
